# Helminth infection induces RELMα-dependent adipose tissue transcriptional reprogramming and protection against diet-induced obesity

**DOI:** 10.64898/2026.06.04.730169

**Authors:** Jennell Jennett, Rebecca E. Ruggiero-Ruff, Ka Man Lam, Kyle Anesko, Gang Chen, Sarah Midou, Shaokui Ge, Meera G. Nair

## Abstract

Obesity, a rising epidemic in the United States, is causally associated with adipose tissue inflammation, therefore targeting this immune dysfunction offers promising therapeutic avenues. Chronic helminth infection can prevent obesity pathogenesis but there remain opportunities to determine the immune factors that mediate protection and whether long-term protection is associated with changes in the visceral adipose tissue. We optimized a mouse model of western diet-induced obesity and investigated if transient helminth infection was protective through the immunoregulatory protein RELMα. Wild-type (WT) or RELMα knockout (KO) C57BL/6J mice were fed a western diet of high glucose and high fat followed by infection with helminth *Nippostrongylus brasiliensis*, which infects the lung and small intestine but is cleared within two weeks. Infection attenuated weight gain and improved glucose tolerance even after the parasite was expelled in WT but not RELMα deficient mice. This protection was associated with reduced adipocyte hypertrophy in WT mice. Adipose tissue bulk RNA sequencing and digital cell quantification indicated that RELMα promoted enrichment of eosinophils and M2 macrophages and upregulated pathways associated with fatty acid oxidation, mitochondrial function, and thermogenesis, whereas RELMα deficiency led to pro-inflammatory, fibrotic, and lipid accumulation transcriptional profiles. Specific genes that were changed with infection and RELMα deficiency included collagen and serpin genes associated with tissue remodeling and fibrosis, and changes within the adipose tissue was confirmed by immunofluorescent staining. Together, these findings establish that helminth-induced RELMα critically protects against western diet–driven obesity and associated adipose transcriptional reprogramming and tissue remodeling.

## Introduction

Obesity is one of the fastest growing public health concerns in the United States and is projected to affect nearly half of adults by 2030^1,2^. Beyond excess adiposity, obesity is increasingly recognized as a chronic inflammatory disease that contributes to type 2 diabetes, cardiovascular disease, and premature mortality^3,4^. At the immunological level, obesity is characterized by profound remodeling of the adipose tissue microenvironment. Lean adipose tissue maintains a type 2 immune environment, enriched in eosinophils and alternatively activated/M2 macrophages, which promotes adipocyte homeostasis, limits fibrosis, and preserves insulin sensitivity. In contrast, obesity drives a type 1 inflammatory shift marked by infiltration of neutrophils and classically activated/M1 macrophages, increased production of TNFα, IL1β, and IFNγ, and enhanced extracellular matrix deposition, leading to adipocyte hypertrophy, fibrosis, and insulin resistance^3–14^.

Given the inflammatory basis of obesity, immunomodulatory strategies have gained interest as potential therapies^15^. Helminth infections represent a natural and potent inducer of type 2 immunity to protect from metabolic disease^11,16–18^. Epidemiological studies indicate that helminth-exposed populations exhibit reduced prevalence of metabolic disease, and experimental models demonstrate that helminth infection attenuates diet-induced weight gain, improves glucose tolerance, reduces hepatic steatosis, and reshapes adipose tissue immunity^11,16,18–20^. Mechanistically, helminths promote polarization of M2 Macs, expansion of eosinophils, and production of type 2 cytokines that collectively suppress obesity-associated inflammation and alter adipose tissue remodeling^11,16,18,21,22^.

One candidate effector linking helminth-induced type 2 immunity to metabolic protection is Resistin-Like Molecule α (RELMα), also known as FIZZ1 or HIMF^23^. RELMα is expressed by macrophages following IL-4 and IL-13 signaling induced during helminth infection ^23–25^. It has been implicated in wound healing, extracellular matrix remodeling, angiogenesis, and resolution of inflammation ^25–27^. In diet-induced obesity, CD301b mononuclear phagocyte secretion of RELMα was beneficial in maintaining a positive energy balance ^21^. RELMα expression exhibits sexual dimorphism, with female mice displaying higher basal RELMα expression and resistance to diet-induced obesity compared to males. In high fat diet-induced obesity, loss of RELMα accelerated weight gain in females, whereas reconstitution of RELMα restored metabolic protection ^22^. These findings suggest that RELMα serves as a molecular mediator of type 2 immune-driven metabolic regulation ^21,22,24,25^.

Despite these observations, critical gaps remain. It is unclear whether helminth-induced metabolic protection persists beyond the acute phase of infection and whether RELMα is required to sustain these effects. In addition, most prior studies have focused on high fat diet models that do not fully recapitulate the high fat and high sugar composition of human western diets. Whether RELMα confers protection in the context of western diet-induced obesity remains unknown. To address these gaps, we evaluated whether RELMα is necessary for helminth-driven metabolic protection during western diet–induced obesity.

## Results

### Western diet–induced weight gain is influenced by sex and RELMα genotype

Sex differences in diet-induced obesity have been reported previously, with WT females demonstrating significant protection from high-fat diet–induced weight gain compared to males, whereas lack of RELMα abrogates this protection^22^. We sought to investigate sexual dimorphism and RELMα expression using a translationally relevant WD model, incorporating high fat and high sugar to better mimic the human dietary patterns responsible for diet-induced obesity. Male and female WT or KO mice were placed on Control Diet (CD) or Western Diet (WD), and body weight trajectories were tracked for up to 15 weeks (Fig. 1A). Body weight trajectories revealed sex- and genotype-dependent responses to WD feeding (Fig. 1B). To determine interactions between diet and sex, linear mixed models were conducted with datasets from two separate experiments (Table 1). A significant sex × diet interaction was detected, indicating that WD-associated weight gain differed between males and females (Table 1, p = 0.0066). When analyzed separately by sex, WD significantly increased body weight in males compared to CD controls (Table 1, p = 0.0075). Both WT and KO males developed WD-associated weight gain, allowing us to test whether helminth-mediated attenuation of obesity is RELMα-dependent. In contrast, female mice showed a genotype-dependent response to WD, with KO females gaining weight on WD while WT females remained largely protected from WD-induced weight gain.

**Figure 1.**
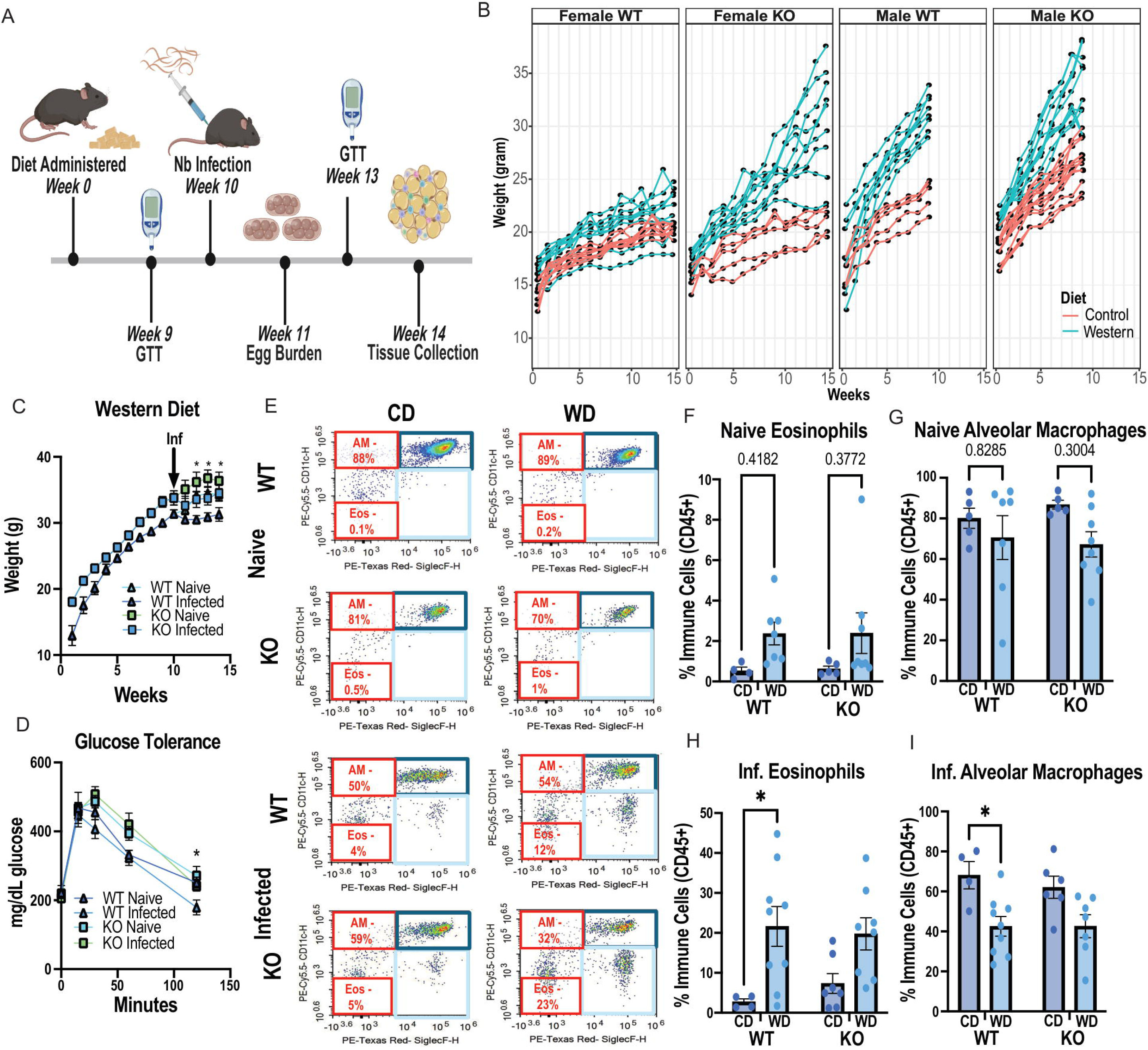
RELMα is required for helminth-mediated attenuation of western diet–induced metabolic dysfunction. **(A)** Experimental timeline. **(B)** Body weight trajectories of male and female WT and KO mice during diet exposure prior to infection. **(C)** Body weight trajectories of male mice before and after *Nippostrongylus brasiliensis* infection at week 10 (arrow). **(D)** Glucose tolerance testing three weeks post-infection, in male mice. **(E)** Representative BALF flow cytometry plots from male mice for alveolar macrophages (CD45 CD11c Siglec-F□) and eosinophils (CD45□ CD11c Siglec-F□). **(F–I)** Frequency of BALF eosinophils (F) and alveolar macrophages (G) among CD45 cells in naïve and infected male mice under CD and WD conditions. Data are presented as mean ± SEM and representative of 2 independent experiments with n = 3–5 mice per group per experiment. *p<0.05 by two-way ANOVA with Tukey’s multiple comparisons (F-I), endpoint analysis (C), and AUC analysis (D).

**Table 1.** Sex Differences in Weight.

| Group | Interaction Terms | Effect size | 95% CI | *p* value |
| --- | --- | --- | --- | --- |
| Male and Female | Sex × Diet | 2 | 0.58 to 3.43 | 0.0066 |
| Male only | Diet (Western vs Control) | 1.92 | 0.55 to 3.28 | 0.0075 |
| Female only | Diet (Western vs Control) | 1.8 | 0.39 to 3.22 | 0.014 |

Because this study aimed to determine whether helminth infection attenuates established obesity in a RELMα-dependent manner, subsequent infection studies focused on male mice.

### Western diet-induced obesity is improved with helminth infection in a RELMα-dependent manner and is associated with increased eosinophil responses but higher parasite burden

The effect of helminth infection on body weight, glucose tolerance, and the immune response was examined in WD-fed WT or KO male mice. Following infection with *Nippostrongylus brasiliensis*, WD-fed WT males had a significant reduction in weight gain compared to naïve WD controls (Fig. 1C). In contrast, KO males were not protected from weight gain post-infection. Linear mixed models identified a significant diet × infection interaction for body weight (Table 2, p < 0.0001). The interaction between genotype and infection was also significant (Table 2, p = 0.007), confirming infection-associated effects that differed between WT and KO males. A Glucose Tolerance Test (GTT) measured three weeks post-infection indicated genotype-dependent differences in glucose tolerance following helminth infection (Fig. 1D). Linear mixed models identified a significant diet × infection interaction for GTT (Table 3, p = 0.04). A significant genotype × diet interaction was also detected (Table 3, p = 0.004), indicating genotype-dependent differences in glucose tolerance under WD conditions. The effect of diet and helminth infection on the immune response was assessed in the bronchio-alveolar lavage (BAL), with quantification of eosinophils, an indicator of a persistent type 2 immune response, and alveolar macrophages, the dominant cell subset in the BAL (Fig. 1E, Fig. S1).

**Table 2.** Infection Effects on Weight in Males.

| Interaction Terms | Associated Weight Change | 95% CI | *p* value |
| --- | --- | --- | --- |
| Genotype (KO vs. WT) | 0.99 | 0.23 – 1.74 | 0.01 |
| Diet (Western vs. Control) | 2.26 | 1.23 – 3.30 | < 0.0001 |
| Infection Status: (Infected vs. Naive) | 1.75 | 0.68 ~ 2.82 | 0.002 |
| Diet × Infection | −2.69 | −3.63 – 1.76 | < 0.0001 |
| Genotype: KO × Infection Status: Infected | 1.30 | −2.21 – 0.38 | 0.007 |
| Weight at Infection Day 0 | 0.95 | 0.87 – 1.04 | < 0.0001 |

**Table 3.** Infection Effects on Glucose Tolerance Test in Males.

| Variable | Effect Size (mg/dL) | 95% CI | *p* value |
| --- | --- | --- | --- |
| Genotype (KO vs. WT) | 0.85 | 0.74 – 0.98 | 0.02 |
| Diet (Western vs. Control) | 1.22 | 1.03 – 1.44 | 0.02 |
| Log scaled Glucose (mg/dL) | 1.75 | 1.41 – 2.17 | <0.0001 |
| Infection Status: (Infected vs. Naive) | 1.06 | 0.94-1.02 | 0.32 |
| Genotype: KO × Diet: Western | 1.27 | 1.09 – 1.49 | 0.004 |
| **Diet: Western Diet ×**<br>**Infection:Infected** | 0.85 | 0.73 – 0.99 | 0.04 |

Under steady-state naïve conditions, there were no significant changes in eosinophil or macrophage frequencies in response to WD or RELMα deficiency (Fig. 1F, G). At three weeks post-infection, eosinophil frequencies were significantly increased in WD compared to CD-fed WT mice, in parallel with a decrease in alveolar macrophages (Fig. 1H, I). In contrast, no significant differences were observed in KO mice. This sustained increase in eosinophils suggests a delayed return to homeostasis in WD-fed mice even after parasite clearance, highlighting an enhanced type 2 response that is consistent with the metabolic improvements seen in helminth-infected WT mice.

While helminth infection conferred metabolic and immunological benefits, the severity of pre-existing obesity impacted parasite clearance. Parasite egg burden at day 7 post-infection indicated a significant increase in fecal eggs with WD compared to CD (Fig. S2; Table 4, p = 0.04), though there were no significant differences between WT and KO mice (Table 4, p = 0.68). Body weight at the time of infection was positively associated with egg burden (Table 4, p = 0.01), and glucose intolerance was also associated with egg burden (Table 4, p = 0.03), suggesting that mice with greater metabolic dysfunction exhibited higher parasite burden. Together, these data demonstrate that helminth infection mitigates WD-induced obesity and is associated with increased eosinophil responses. Helminth infection additionally improved glucose tolerance in WT mice, but this effect was not observed in KO mice, suggesting RELMα-dependent protection from WD-associated glucose intolerance. While these metabolic improvements occurred in WT mice, the severity of pre-existing obesity and glucose intolerance was a predictor for higher parasite egg burden across genotypes.

**Table 4.** Diet Effects on Parasite Egg Burden in Males.

| **Variable** | **Effect Size**<br>**(eggs/gram)** | **95 % CI** | ***p* value** |
| --- | --- | --- | --- |
| **Genotype (KO vs. WT)** | 0.76 | 0.19 – 2.99 | 0.68 |
| **Diet (Western vs. Control)** | 4.36 | 1.10 – 17.22 | 0.04 |
| **GTT per unit** | 0.001 | 0.00 – 0.08 | 0.03 |
| **Weight** | 1.30 | 1.06 – 1.59 | 0.01 |

### Helminth infection reduces adipocyte size and is associated with transcriptional changes in the adipose tissue immune responses

Changes in the visceral adipose tissue with RELMα deficiency and in response to helminth infection was examined in Hematoxylin and Eosin (H&E) stained perigonadal adipose tissue sections of WD-fed mice (Fig. 2A). Quantitative analysis revealed that while naive WT and both naive and infected KO males exhibited comparable adipocyte size, helminth infection significantly reduced adipocyte size exclusively in WT males (Fig. 2B). To determine if RELMα-dependent reduction in adipocyte size was linked to local immunological shifts, we evaluated transcriptional changes in the adipose tissue by bulk RNA sequencing data and digital cell quantification (DCQ) ^28,29^. While WT and KO samples were broadly similar across most immune populations under naive conditions, WT mice consistently maintained a higher estimated abundance of eosinophils and M2 macrophages across both naive and infected states. In contrast, KO samples exhibited a baseline preference in M1 macrophages, which evolved into a broader type 1 pro-inflammatory microenvironment following infection. Specifically, infection reduced the estimated abundance of memory B cells, naive CD4+ T cells, regulatory T cells, and conventional dendritic cells in both genotypes, with these populations reaching their lowest levels in WT infected samples. Conversely, estimated monocyte and neutrophil abundance increased with infection; however, these pro-inflammatory populations were more abundant in KO infected samples compared to WT controls. Analysis of differentially expressed genes expression (DEG) (log2 fold change ≥ 0.25 and adjusted p ≤ 0.05) revealed that there were few DEGs between WT and KO mice under naïve conditions (22 DEG), which were mostly upregulated in KO mice (21 DEG) (Fig. 3B, Fig S3). In contrast, infection drove a 7.4-fold increase in DEGs between WT and KO mice demonstrating a profound genotype-dependent shift during infection. Within genotypes, infection induced extensive transcriptional changes in WT mice (433 DEG) (Fig. 3C). In contrast, infection induced 33.3-fold fewer DEGs in KO mice.

**Figure 2.**
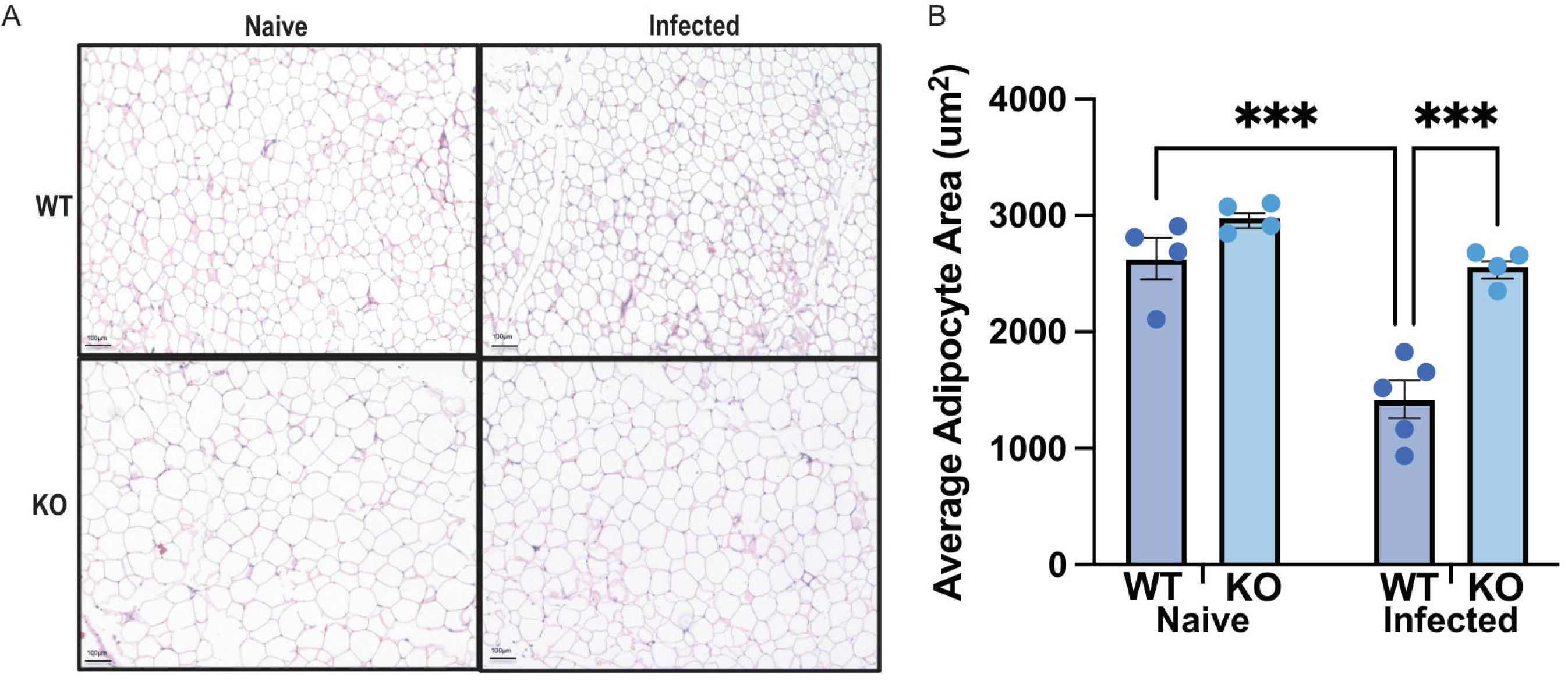
RELMα is required for infection-associated reduction in adipocyte size. **(A)** Representative hematoxylin and eosin (H&E) stained visceral adipose tissue sections from WT and KO naïve and *Nippostrongylus brasiliensis*-infected male mice. Scale bar, 100 µm. **(B)** Average adipocyte area (µm²). Data are presented as individual animals with group means ± SEM. Statistical analysis was performed using two-way ANOVA with Tukey’s multiple comparisons test. Data are representative of 2 independent experiments with n = 3–5 mice per group per experiment (***p < 0.001).

**Figure 3.**
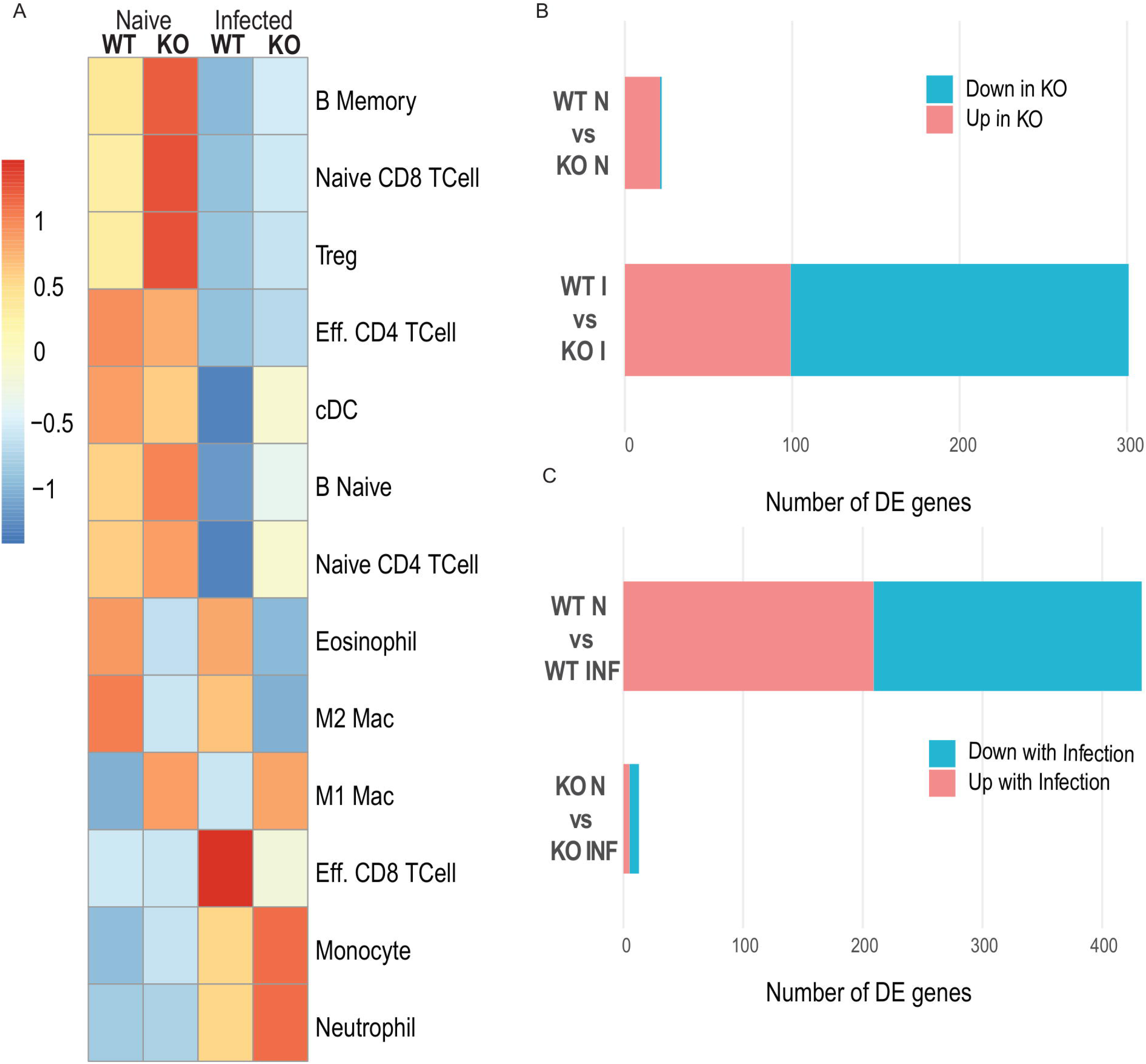
RELMα deficiency alters infection-associated immune composition and transcriptional responses in adipose tissue. **(A)** Digital cell quantification (DCQ) analysis of bulk adipose RNA sequencing data estimating relative immune cell abundance in WT and KO mice under naïve and infected conditions. Heatmap represents relative enrichment scores across immune populations. **(B)** Differential gene expression between WT and KO adipose tissue under naïve and infected conditions. **(C)** Within-genotype differential gene expression comparing naïve and infected conditions. DEG were defined as absolute log2 fold change ≥ 0.25 with adjusted p value ≤ 0.05. Representative of 1 experiment, n=3-4 per group.

Together, these data indicate that helminth infection induces substantial transcriptional changes in the visceral adipose tissue that are dependent on RELMα. Through digital cell quantification, genotype and infection-associated changes in immune cell subsets included increased eosinophils and M2 macrophages in WT mice while M1 macrophages, monocytes and neutrophils were higher in KO mice.

### Helminth infection decreases adipose extracellular matrix and tissue remodeling programs and upregulates cellular metabolism pathways in a RELMα-dependent manner

Comparison of the top 30 DEG between adipose tissue from naïve and infected WT mice indicated genes associated with extracellular matrix organization and metabolic regulation (Fig. 4A). Transcripts involved in ECM composition and tissue remodeling (*Col12a1, Fgf18, Fbln2, Hmcn2)* were reduced in WT infected samples relative to WT naive controls, whereas genes linked to lipid and small molecule metabolism (*Tmem88b, Cyp2e1, Nat8l*) were increased ^5,6,12,30–44^. This was consistent with Gene Ontology (GO) biological process analysis indicating downregulation of ECM organization, collagen metabolic processes, connective tissue development, and wound healing pathways with helminth infection ^5,6,12,26,30,45–47^(Fig. 4B).

**Figure 4.**
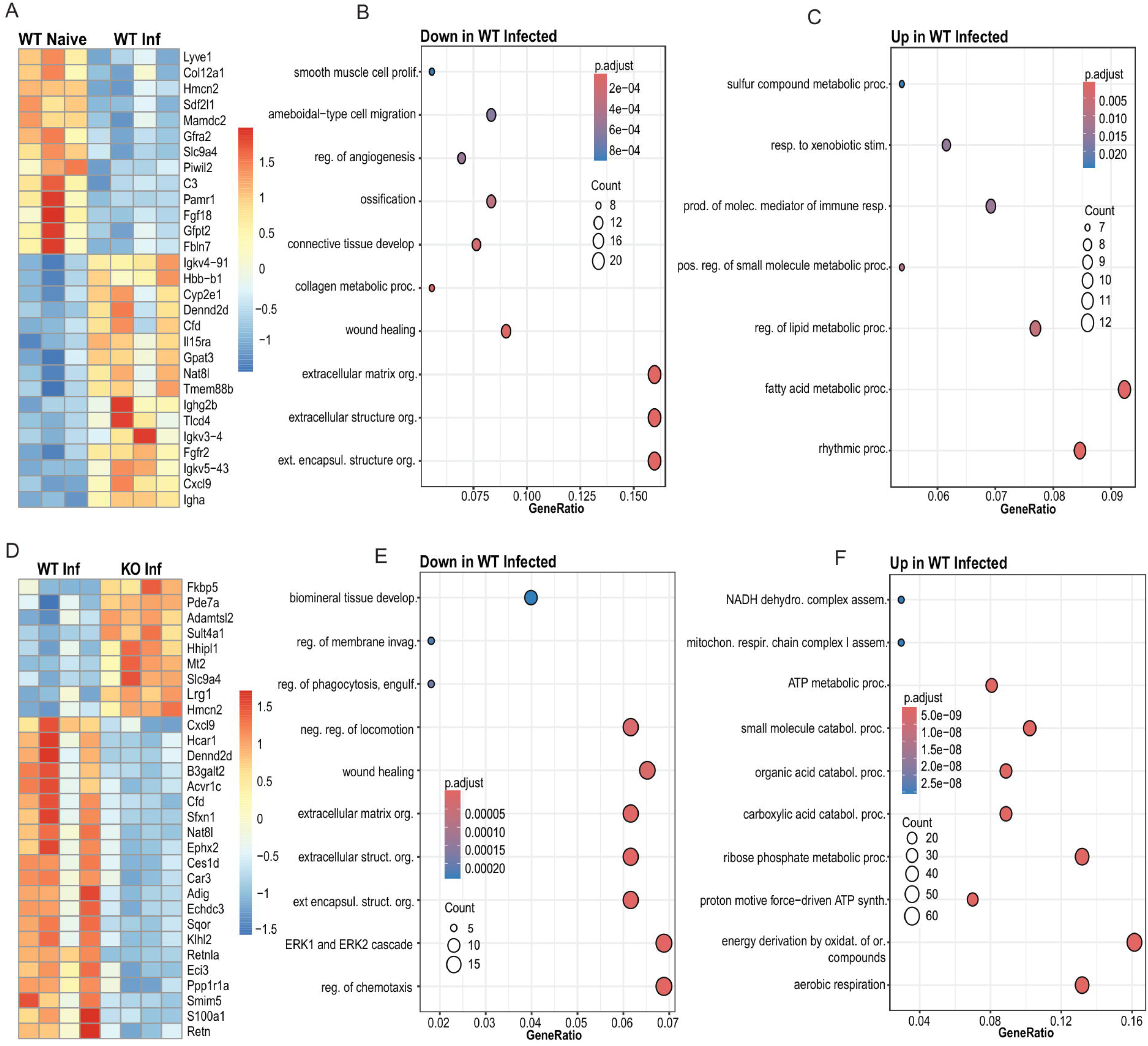
**Infection-associated adipose tissue suppression of extracellular matrix genes and induction of oxidative metabolism are RELM**α**-dependent. (A-C)** Heatmap of differential gene expression analysis of WT infected versus WT naïve adipose tissue (A) and Gene Ontology (GO) Biological Process enrichment for downregulated (B) upregulated pathways (C). **(D-F)** Heatmap of differential gene expression analysis of WT infected versus KO infected adipose tissue (D) and GO Biological Process enrichment analysis of downregulated (E) or upregulated pathways (F) reduced in WT infected relative to KO infected adipose tissue. Genes selected met the criteria of absolute log2 fold change ≥ 0.25 and padj ≤ 0.05 are shown. Representative of 1 experiment, n=3-4 per group per experiment.

Genes upregulated in WT infected samples were enriched for fatty acid and lipid metabolic processes and oxidative metabolism, consistent with improved metabolic function with helminth infection ^11,16,18–20,48^ (Fig 4C). Comparison of WT infected and KO infected samples revealed genotype-dependent transcriptional differences (Fig. 4D). The top genes expressed in KO mice were involved with stress and metabolic signaling (*Fkbp5, Mt2, Pde7a*) ^35,49–51^. Conversely, there were more genes increased in WT infected relative to KO infected samples, including genes associated with adipocyte maintenance and function (*Cfd*, *Adig*, *Hcar1*) and metabolic activity (*Eci3, Echdc3*) ^17,52–59^. Gene ontology analysis indicated downregulated tissue remodeling pathways in WT mice (eg. extracellular matrix organization, wound healing) (Fig. 4E), while pathways involved in aerobic respiration, mitochondrial respiratory chain complex assembly, and ATP synthesis were upregulated (Fig. 4F), aligned with improved metabolic activity in WT mice. There were few genes that were differentially expressed between WT and KO mice under naïve conditions (Fig. S3A), or between KO naïve and infected conditions^60,61^(Fig. S3B). This suggested that RELMα deficiency did not impact in the adipose tissue under homeostatic conditions but that it is a driver of transcriptional reprogramming following helminth infection, with differentially expressed genes that affect obesity and glucose tolerance outcomes ^21–25,52^.

To define these RELMα-dependent DEGs that are associated with helminth-induced protection from obesity, we analyzed genes significantly altered in WT infected versus WT naive samples but unchanged in KO mice (Fig. 5A). We identified 26 genes that could be categorized into three main functions: immune, structural and metabolic. Immune genes showed a similar distribution of upregulated and downregulated genes with infection in WT mice. Upregulated genes were associated with T cell recruitment (*Cxcl9*, *Il15ra*, *Cd4*) but surprisingly, genes associated with M2 macrophages, which are associated with protective outcomes in obesity, were downregulated (*Il4ra*, *Cd163*) ^11,16,18,20,21,24,48,62–67^. All genes with structural functions were significantly downregulated with infection in WT mice but not KO mice (*Col14a1, Fbln7, Fbln2, Serpinh1, Serpinf1, Col6a1*) ^5,6,12,30,68–73^. GO analysis of these WT-specific downregulated transcripts confirmed enrichment for ECM organization, collagen metabolic processes, connective tissue development, and wound healing ^5,6,12,26,30^ (Fig. 5B). Within metabolic activity-associated genes, only *Cdkn1a*, which is a cell cycle inhibitor induced by cellular stress and DNA damage, was significantly downregulated in infected mice. In contrast, 7 genes were significantly upregulated in infected WT but not KO mice, many of which were associated with lipid metabolism ^31–34,38,61,74,75^ (*Cyp2e1, Cyp2f2, Pdk2*). Consistent with this, GO analysis of WT-specific upregulated genes showed enrichment in regulation of lipid and fatty acid metabolic processes and triglyceride biosynthesis (Fig. 5C), which were not observed in the KO adipose tissue ^60^.

**Figure 5.**
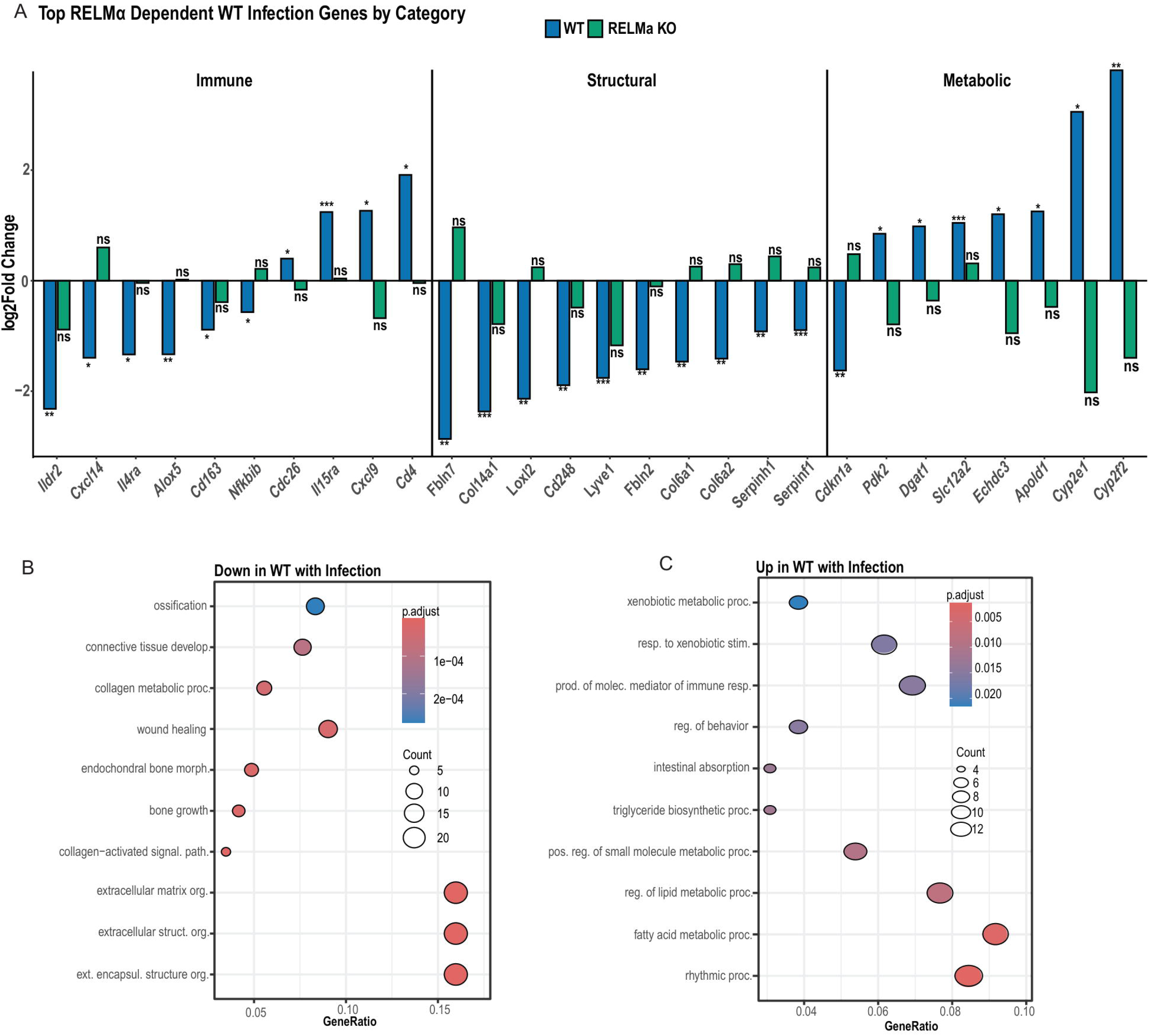
RELMα governs WT-specific transcriptional responses to infection in adipose tissue. **(A)** Top RELMα-dependent infection-associated genes grouped by functional category (Immune, Structural, Metabolic). RELMα-dependent genes were defined as transcripts significantly altered in WT infected relative to WT naïve adipose tissue (DESeq2; absolute log2 fold change ≥ 0.25, padj ≤ 0.05) and not significantly changed in KO mice. Bars represent log2 fold change with infection in WT (blue) and KO (green). Significance for each genotype is indicated as shown (*padj < 0.05; **padj < 0.01; ***padj < 0.001; ns, not significant). **(B-C)** GO Biological Process enrichment analysis of RELMα-dependent genes downregulated (B) or upregulated (C) in WT mice with infection. Representative of 1 experiment, n=3-4 per group per experiment.

Taken together, these transcriptomic data provide insight into the protective RELMα-dependent pathways in the visceral adipose tissue that are induced by helminth infection. First, it appears that helminth infection coordinates a RELMα-dependent structural transition in adipose tissue from a rigid structural focus to an active metabolic state. This process involved the simultaneous downregulation of extracellular matrix and tissue remodeling programs—such as *Col12a1* and *Fgf18*—while genes involved in fatty acid metabolism and mitochondrial respiratory chain assembly were upregulated ^9,12,40,41,44^. In contrast, the adipose tissue in RELMα-deficient mice did not undergo these metabolic and remodeling pathways and instead exhibited cellular stress and metabolic arrest characterized by high levels of *Fkbp5* and *Mt2*^49,50,76^. Overall, these findings suggest that RELMα is a critical molecular switch that allows infection to trigger beneficial adipose reprogramming for protection against obesity.

### Changes in tissue remodeling with helminth infection is associated with differential expression of collagen and serpin proteins

Based on the significant changes in structural genes in WT infected, we investigated whether these changes are key tissue structure-associated proteins by immunofluorescent staining of adipose tissue sections (Fig 6). Collagen VI staining was significantly reduced in infected WT mice relative to naïve WT controls, whereas this reduction was not observed in KO mice (Fig. 6A). In addition, infected KO mice exhibited significantly greater collagen VI deposition than infected WT mice. These findings are consistent with the RNA sequencing data and support a model in which RELMα is required for infection-associated suppression of adipose extracellular matrix deposition. We next assessed Serpine1, another remodeling associated marker identified in our transcriptomic analysis. Serpine1 staining differed significantly by genotype, with KO mice exhibiting increased signal relative to WT mice under both naïve and infected conditions, although infection did not significantly alter Serpine1 deposition (Fig. 6B). Thus, unlike collagen VI, serpine1 appeared to be influenced primarily by genotype rather than infection status. Lyve1, expressed by endothelial cells and perivascular macrophages was among the transcripts altered in our RNA sequencing analysis, however, Lyve1 protein expression did not significantly differ by genotype or infection status (Fig. 6C).

**Figure 6.**
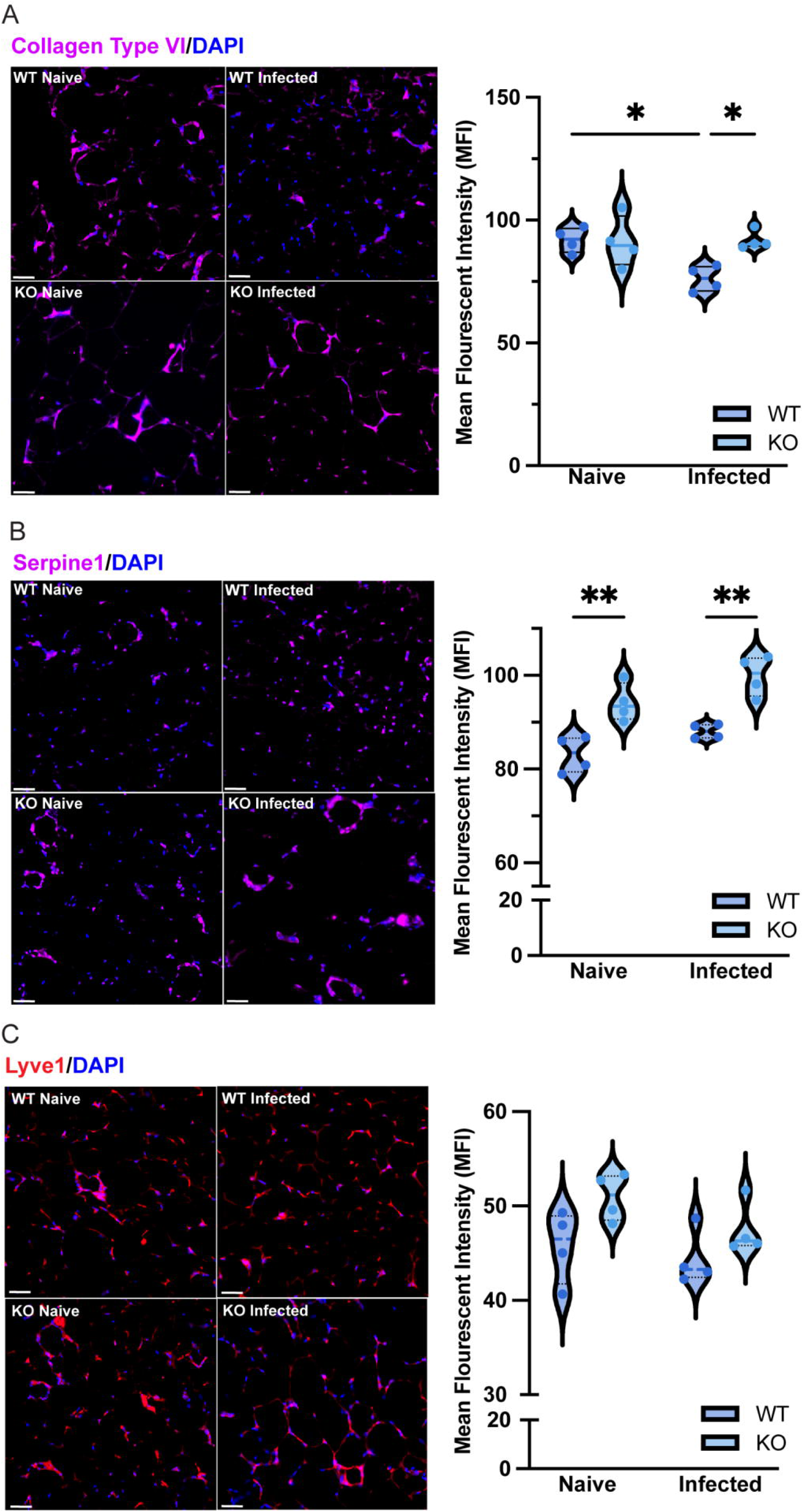
RELMα regulates adipose tissue extracellular matrix associated protein deposition in adipose tissue. Representative immunofluorescence images and quantification of Collagen VI (A), Serpine1 (B), Lyve1 (C) in visceral adipose tissue from WT and KO male mice under naïve and infected conditions. Nuclei were counterstained with DAPI. Scale bar = 20uM. Data are represented as individual animals with group means. Statistical analysis was performed by two-way ANOVA with Tukey’s multiple comparisons test. Data are representative of 2 independent experiments, n = 4 mice per group, n=10 photos per mouse (*p < 0.05; **p< 0.01).

Together, these data confirm that RELMα is required for helminth-induced reduction of Collagen VI, a key structural component of the adipose extracellular matrix. The suppression of collagen deposition during helminth infection in WT but not KO mice may allow for beneficial adipose tissue remodeling for increased metabolic activity to reduce obesity. Instead, adipose tissue from KO mice had elevated levels of Serpine1 in both naïve and infected conditions, suggesting a baseline defect in tissue remodeling capacity independent of infection ^12,43,70,77^.

Combined, these transcriptomic and protein data points to the intriguing importance of adipose tissue structure for optimal whole body metabolic outcomes, with the long-term protective role of RELMα-mediated tissue remodeling in response to a transient helminth infection.

## Discussion

The goal of this study was to determine whether transient helminth infection protects against western diet-induced obesity after parasite clearance and whether this protection requires RELMα. We found that infection with helminth *Nippostrongylus brasiliensis* attenuated weight gain, improved glucose tolerance, and limited adipocyte size in western diet-fed WT male mice, whereas these protective effects were absent in KO animals. Together, these findings identify RELMα as a required mediator of helminth-associated protection from metabolic dysfunction and extend prior work showing that helminth infection can improve obesity-associated outcomes through type 2 immune pathways and M2 macrophage^5,11,15,16,18,20,24^.

An important feature of this model is that the metabolic benefit of infection was sustained after the acute phase of infection, indicating that transient helminth exposure can alter host metabolic responses beyond the period of active parasite burden. Prior studies have shown that RELMα contributes to tissue repair and post-infection remodeling in helminth infection settings, supporting the broader concept that RELMα-dependent pathways can shape tissue adaptation after infection resolution ^24–27^. Although those studies were performed in the lung rather than adipose tissue, our findings raise the possibility that helminth-induced RELMα similarly contributes to sustained adipose tissue remodeling after infection, and that these structural changes are critical for adipose organ metabolic activity for improved obesity outcomes. These results also extend on previous work on RELMα in high fat diet-induced obesity but investigate the effects of western diet and helminth infection. Prior studies showed that RELMα regulates adipose macrophages and eosinophils and contributes to sex-specific protection from high fat diet-induced obesity, with female mice showing resistance to obesity that is lost in the absence of RELMα ^22,25^. Following western diet, however, WT females were resistant to weight gain, whereas KO females lost this protection. This indicates that sex differences are more pronounced with western diet than high fat diet DCQ analysis helped clarify the cellular context of these transcriptional changes. WT adipose tissue was characterized by higher estimated eosinophil and M2 macrophage abundance, whereas KO adipose tissue had relatively increased M1 macrophages and monocytes. These findings are consistent with a model in which RELMα supports a type 2 skewed adipose immune environment that is more permissive to metabolic homeostasis and less prone to chronic inflammatory remodeling. This interpretation is supported by prior work showing that RELMα induction is closely linked to IL-4 driven macrophage polarization and helminth associated macrophage programming ^16,18,22,24,25,27,78^. These results suggest that RELMα is not only associated with type 2 activation but may also help regulate macrophage function and adipose tissue metabolic homeostasis.

Another finding was that consumption of a WD increased parasite egg burden. In addition, post-infection glucose levels and body weight at the time of infection were positively associated with egg burden. These observations suggest that host metabolic state may influence parasite fitness, host defense, or both. Although RELMα genotype alone was not significantly associated with egg burden, the association between worsened metabolic status and increased egg burden raises the possibility that obesity and over-nutrition create a more permissive environment for parasite development or reproduction. More broadly, this finding supports the concept that the relationship between helminth infection and metabolism is bidirectional, with immune and metabolic state shaping not only host pathology but also parasite associated outcomes ^15,20,48,79,80^. WD also had significant effects on the pulmonary tissue microenvironment. During infection, WD fed mice had higher BALF eosinophils than CD fed mice, with a similar trend present in naïve animals. This suggests that WD promotes a more eosinophilic state in the lung that becomes more apparent during helminth infection. This is notable because obesity is increasingly recognized as a modifier of airway inflammation, and prior studies in allergic airway models have similarly shown that high fat diet or obesity can enhance eosinophilic lung inflammation and augment type 2 associated airway responses^13,27,81–86^. Thus, although our model uses helminth infection, the BALF data are consistent with the broader idea that overnutrition can bias the lung toward a stronger type 2 associated inflammatory state.

Additionally, WD increased egg burden, yet this occurred in conjunction with increased pulmonary eosinophilia rather than its absence. This suggests that overnutrition does not blunt the host response to infection but may instead alter its quality. In that setting, inflammatory features of the response are maintained or increased without corresponding to improvement in parasite associated outcomes.

Histologic and transcriptomic data further supports a role for RELMα in regulating adipose tissue remodeling during obesity. Infection reduced adipocyte size in WT mice, whereas this effect was absent in KO mice. In parallel, infected WT adipose tissue showed reduced expression of extracellular matrix and structural remodeling programs, including pathways related to collagen organization, connective tissue development, and wound healing. These findings are notable because obesity is strongly associated with extracellular matrix remodeling, collagen deposition, and fibrosis, all of which contribute to adipose dysfunction and insulin resistance ^5,6,9,10,12,35,37,39,49,70,87–89^. The coordinated reduction in matrix associated pathways in infected WT animals therefore aligns closely with the morphologic finding of smaller adipocytes and supports the interpretation that RELMα helps limit maladaptive tissue remodeling in obese adipose tissue. At the same time, infected WT adipose tissue showed enrichment of pathways related to fatty acid metabolism, small molecule metabolism, aerobic respiration, and mitochondrial function, whereas these changes were not equally enriched in KO mice. These findings suggest that RELMα promotes a more metabolically favorable transcriptional response in adipose tissue following helminth infection. This interpretation is consistent with prior work showing that eosinophils, type 2 cytokines, and M2 macrophages contribute to metabolically protective adipose remodeling and beige fat associated pathways ^5,11,16,18,21,22,24,65,85^. However, our study did not directly measure thermogenesis, mitochondrial respiration, or energy expenditure and future studies are warranted to study infection-related increase in thermogenic function and the influence of RELMα.

The findings reported here further indicate that helminth-induced changes in adipose tissue structure require RELMα. When WT-specific infection-responsive genes were examined, infection reduced a coordinated set of extracellular matrix and structural genes while inducing genes involved in lipid metabolism and other metabolic processes. These same transcriptional shifts were not seen in KO mice. Thus, helminth infection alone was not sufficient to improve the adipose tissue state. Instead, the beneficial adipose response to infection appeared to depend on RELMα. This suggests that RELMα is required for adipose tissue to interpret infection induced immune signals in a way that limits persistence of a matrix rich, obesity associated phenotype and permits a more metabolically adaptive response. The reduction in the expression of genes such as Col6a1, Fbln1, and Fbln2 is consistent with decreased fibrotic and structural remodeling, whereas induction of Dgat1 and Echdc3 suggests a shift toward lipid handling and metabolic adaptation ^5,6,12,17,30,33,90–93^. These findings parallel our previously described phenotype in WT mice, where infection improved metabolic outcomes and reduced adipocyte size, and further support the idea that without RELMα the adipose tissue microenvironment is not similarly benefited by infection.

Adipose tissue immunofluorescent staining indicated that Collagen VI was among the extracellular matrix associated genes reduced in WT adipose tissue after infection but not in KO mice, and immunostaining showed a similar pattern at the protein level. Collagen VI deposition was reduced in infected WT mice, whereas this reduction was not observed in infected KO mice. Infected KO mice also retained greater collagen VI staining than infected WT mice. This finding is consistent with prior work showing that collagen VI is increased in obese adipose tissue and is associated with fibrosis, macrophage accumulation, and insulin resistance^5,6,9,10,12,37,39,44,70,89^.

Experimental studies have further shown that collagen VI contributes to metabolic dysfunction in obesity, and that endotrophin, a cleavage product of collagen VI, promotes adipose fibrosis and systemic insulin resistance^30^. In this context, the reduction in collagen VI deposition in infected WT mice supports the interpretation that RELMα contributes to limiting fibrotic remodeling in adipose tissue after infection. The mechanism by which RELMα regulates this response remains unclear. One possibility is that RELMα alters the adipose immune environment in a way that limits the persistence of a matrix-rich inflammatory state. This interpretation is consistent with our transcriptional data, which identified RELMα dependent changes in structural and immune pathways, and with prior work supporting a role for RELMα in maintaining metabolic homeostasis and regulating tissue remodeling ^21–27^. Although our data does not distinguish whether RELMα acts directly on matrix-producing stromal cells or indirectly through macrophages and other immune populations, it supports a model in which RELMα is required for helminth infection associated reduction in extracellular matrix deposition in obese adipose tissue. Serpine1 followed a different pattern. Serpine1 staining was increased in KO mice under both naïve and infected conditions, whereas infection itself did not significantly alter deposition. Serpine1 has been linked to adipose inflammation, macrophage recruitment, and metabolic dysfunction in obesity, suggesting that RELMα deficiency may favor a broader remodeling associated tissue state rather than a discrete infection dependent change in this pathway^69,70,89^.

There remain limitations in this study. First, *Nippostrongylus brasiliensis* is a live infection model, and it remains unclear whether the safer therapeutic option of helminth-derived products would induce similar RELMα dependent metabolic protection. Second, our adipose sequencing was performed on bulk tissue, which limits cellular resolution and prevents direct assignment of transcriptional changes to adipocytes, macrophages, eosinophils, or stromal populations. Third, although pathway analysis supported oxidative and mitochondrial transcriptional programs in infected WT adipose tissue, functional studies will be required to determine whether these changes translate into altered adipose respiration, substrate utilization, or energy expenditure^11,17,60,94^.

In summary, these studies identify a previously unrecognized role for helminth induced RELMα in shaping adipose tissue responses to western diet-induced obesity. Our findings support a model in which RELMα promotes infection associated metabolic protection by limiting adipocyte hypertrophy, restraining extracellular matrix remodeling, and supporting a more oxidative and type 2 associated adipose tissue environment^4,9–11,16,18,22,27,58^.

## Materials and Methods

### Mouse studies

All experiments were conducted in accordance with protocols approved by the Animal Care and Use Committee at the University of California, Riverside (AUP#48). RELMα KO mice, generated as previously described (22), and C57BL6/J WT controls were bred in the vivarium and maintained under a 12-hr light, 12-hr dark cycle. Six- to eight-week-old mice were provided ad libitum access to Control Diet (Research Diets; Cat D14042701, 10% kcal from fat, 73% kcal from carbohydrate, 0% from sucrose) or Western Diet (Research Diets; Cat D12079B, 17% kcal protein, 40% kcal fat, 43% kcal carbohydrate. 29% kcal from sucrose) beginning at week 0 and maintained on diet for the duration of the study. Body weight was recorded weekly during the pre-infection period, and daily after infection with *Nippostrongylus brasiliensis* at week 10 of diet administration. Mice were briefly anesthetized with isoflurane then infected subcutaneously with 550 *N. brasiliensis* L3 larvae or PBS control. Following infection, mice remained on their respective diets and were monitored through the study endpoint at week 14.

Glucose tolerance tests were performed at 9 weeks following initiation of diet and at 3 weeks post infection. Mice were fasted for 6 hours prior to testing. Fasting blood glucose was measured at time 0 using a glucometer following tail blood collection. Mice then received an intraperitoneal injection of glucose solution (200 g/L) (Gibco) at 10 µL per gram body weight. Blood glucose was measured from tail blood at 15, 30, 60, and 120 minutes following injection.

### Bronchoalveolar lavage fluid collection and flow cytometry

Bronchio-alveolar lavage fluid (BALF) was collected by three lavages of 0.7 mL sterile 1X PBS. Standard staining for flow cytometry included CD16/32 (Biolegend, 5 µg/mL) and rat IgG (Millipore Sigma, 10 µg/mL) followed by staining with surface antibody cocktail (0.5 µg/mL/Ab; Table S1). Data was analyzed using Novoexpress flow cytometry analysis software.

### Adipose tissue imaging

Gonadal adipose tissue was collected from PBS-perfused mice and fixed in 10% NBF for 24 hours at room temperature before being moved to 70% ethanol at 4°C and sent to the Translational Pathology Core Laboratory at UCLA for paraffin embedding and H&E staining of 5 μm sections. H&E-stained visceral adipose tissue sections were imaged under brightfield at 10× magnification using a Keyence microscope, 10–17 non-overlapping fields per animal.

Adipocyte area was quantified using the Adiposoft plugin in Fiji. Images were calibrated at 0.76 μm per pixel, and adipocyte detection parameters were restricted to a minimum diameter of 10 μm and a maximum diameter of 100 μm. ROIs corresponding to individual adipocytes were automatically detected by the software. All ROIs were visually inspected and validated. Mean adipocyte area per animal was calculated from all validated ROIs.

For immunofluorescent staining, tissue sections were dewaxed and rehydrated with standard xylene/ethanol/distilled water washes followed by antigen retrieval by boiling slides in 1× antigen retrieval buffer, pH 6.0 (Abcam) (20 min). Sections were encircled with a hydrophobic barrier pen followed by standard antibody staining. Briefly, tissue was permeabilized in 0.4% Triton X 100 in PBS, blocked in 5% goat serum and 2.5% BSA in PBST (PBS containing 0.05% Tween 20) followed by the Avidin/Biotin Blocking Kit (Vector Laboratories, Cat. SP-2001) according to the manufacturer’s instructions. Sections were incubated overnight at 4°C with primary antibodies (Table S2), diluted in PBST supplemented with 2.5% BSA, washed twice in PBST then incubated with secondary antibodies for 2 hours at RT, and washed twice in PBST, followed by a brief rinse in distilled water. Sections were mounted with Vectashield Vibrance Antifade Mounting Medium with DAPI (Vector Laboratories). Slides were allowed to cure overnight at RT and sealed with clear nail polish prior to imaging. To quantify Collagen VI, Serpine1 and Lyve1 expression, immunofluorescent images were analyzed using an automated batch-processing macro in ImageJ/Fiji (v1.54). To ensure unbiased analysis, ten fields of view were randomly selected for each stain per animal.

Images were converted to 8-bit grayscale and subjected to background subtraction using a rolling ball algorithm (radius = 50 pixels) to remove non-specific noise. An automated Otsu thresholding method was applied to create a binary mask, isolating positive signal from the background. Results are reported as the Mean Gray value to determine the average density of fluorescent intensity.

### Adipose tissue RNA sequencing

Adipose tissue was flash frozen at collection and stored at −80°C until processing. For RNA isolation, approximately 10 to 30 mg of tissue was removed and homogenized in TRIzol reagent (ThermoFisher) (1 mL per ≤50 mg tissue) using a rotor stator electric homogenizer.

Samples were incubated for 5 minutes at RT to permit complete dissociation of nucleoprotein complexes. Phase separation was performed by addition of chloroform (200µL per 1mL TRIzol), followed by centrifugation at 12,000 × g for 15 minutes at 4°C. The aqueous phase was transferred to a fresh tube and subjected to a second chloroform extraction. The resulting aqueous phase was mixed with an equal volume of 70% ethanol and applied to Qiagen™ RNeasy spin columns. Columns were washed twice with RW1 buffer, followed by two washes with RPE buffer. An additional wash with 80% ethanol was performed. RNA was eluted in RNase free water prewarmed to 60°C. RNA concentration and purity were assessed by spectrophotometry, and samples were submitted to Novogene Corporation, Inc for library preparation and RNA sequencing.

Raw sequencing reads from adipose tissue samples were assessed for quality using FastQC^95^ and summarized with MultiQC^96^. Adapter sequences and low-quality bases were trimmed using Trimmomatic^97^. Trimmed reads were aligned to the mouse reference genome using STAR aligner ^98^, and gene-level read counts were generated using FeatureCounts^99^ . Downstream analyses were performed in R. Differential gene expression analysis was conducted using DESeq2^100^. Genes with an adjusted p value (Benjamini–Hochberg correction) ≤ 0.05 and an absolute log2 fold change ≥ 0.25 were considered differentially expressed.

Variance stabilized expression values generated by DESeq2 were used for heatmap visualization. Gene Ontology Biological Process enrichment analysis was performed on differentially expressed gene sets using clusterProfiler^101^ with annotation from org.Mm.eg.db^102^. Digital cell quantification analysis^28,29^ was performed in R using established immune reference signatures to estimate relative immune cell population changes from bulk RNA sequencing data.

### Statistical Analyses

To assess the impact of diet, genotype, infection, and time (weeks pre- and post-infection) on each outcome linear mixed models were employed. These factors and their biologically relevant interactions were treated as fixed effects, while individual identity was included as a random effect. Model results are reported as mean effects with associated 95% confidence intervals (CIs) and p-values (Tables 1-4). Data in figures are presented as mean ± SEM and statistical significance was determined using GraphPad Prism with two-way ANOVA and post-tests as indicated. Results are representative of two independent experiments with n=3-5 per group, apart from the RNA-sequencing experiments which were conducted once with n=3 for the naïve groups and n=4 for the infected groups. P-values <0.05 were considered statistically significant.

## Acknowledgments

We would like to thank the following University of California equipment cores: UC Riverside School of Medicine Research Core for the flow cytometry equipment and biostatistics analysis (somresearch.ucr.edu), UC Riverside High Performance Computing Center for bioinformatics data processing (hpcc.ucr.edu); UC Los Angeles Translational Pathology Core Laboratory for tissue processing. This work was supported by the National Institutes of Health (R01AI153195 to MGN and JJ; R21AI180561, R01AI191470 to MGN), and the California Institute for Regenerative Medicine (TRANSCEND Award# EDUC4-12752 to RER and RAMP Award EDUC5-13636 to KML). The contents of this publication are solely the responsibility of the authors and do not necessarily represent official views of CIRM or other agencies of the State of California.

## Supplemental material

**Supplemental Figure 1. Representative gating strategy for analysis of bronchoalveolar lavage fluid immune cells.**

(A) Representative flow cytometry plots showing the gating strategy used to identify alveolar macrophages and eosinophils in bronchoalveolar lavage fluid (BAL). Events were sequentially gated on cells, singlets, and CD45□ immune cells. Alveolar macrophages were defined as CD11c Siglec-F□ cells and eosinophils as CD11c Siglec-F cells within the CD45 population.

**Supplemental Figure 2. Western diet increases parasite egg burden following infection.**

(A) Quantification of parasite egg burden from WT and RELMα KO male mice maintained on CD or WD for 10 weeks prior to infection with 550 *Nippostrongylus brasiliensis* larvae per mouse. Egg burden was assessed at day 7 post-infection. Data are presented as group means ± SEM. Statistical analysis was performed as described in Methods. Representative of 2 independent experiments, n =3–5 mice per group per experiment.

**Supplemental Figure 3. Differential gene expression analysis of adipose tissue from naïve WT versus naïve KO mice and naïve KO versus infected KO mice.**

(A) Differential gene expression analysis of naïve WT versus naïve KO adipose tissue performed using DESeq2. Genes meeting criteria of absolute log2 fold change ≥ 0.25 and adjusted *p* ≤ 0.05 are shown. (B) Differential gene expression analysis of naïve KO versus infected KO adipose tissue performed using DESeq2. Genes meeting criteria of absolute log2 fold change ≥ 0.25 and adjusted *p* ≤ 0.05 are shown. Representative of 1 experiment, *n* = 3 to 4 mice per group.

**Supplemental Table 1.** BALF Flow Cytometry Panel.

| **Laser** | **Fluorophore** | **Surface** | **Company** | **Cat.** | **Clone** |
| --- | --- | --- | --- | --- | --- |
| **488 nm** | FITC | F4/80 | Biolegend | 123108 | BM8 |
|  | PerCP, 7AAD | 7AAD | Biolegend | 420404 | N/A |
|  | PerCP Cy5.5 | CD115 | Biolegend | 135526 | AFS98 |
|  | PE Cy7 | Ly6C | Invitrogen | 25-5932-82 | HK1.4 |
| **637 nm** | APC | CX3CR1 | Biolegend | 149008 | SA011F11 |
|  | AF700 | MHCII | Invitrogen | 50-169-17 | M5/114.15.2 |
|  | APC Cy7 | CD11b | Biolegend | 101226 | M1/70 |
| **405 nm** | Pacific Blue, BV421 | CD163 | Biolegend | 155309 | S15049I |
|  | AmCyan, BV421 | Ly6G | Biolegend | 127627 | 1A8 |
|  | Pacific Orange, BV570 | CD45 | Biolegend | 103135 | 30-F11 |
|  | Qdot 605, BV605 | CD64 | Biolegend | 139323 | X54-5/7.1 |
|  | BV711 | CD206 | Biolegend | 141727 | C068C2 |
| **561 nm** | PE, Td Tomato | CCR2 | Biolegend | 150610 | SA203G11 |
|  | PE Texas Red, PE Dazzle | SiglecF | Biolegend | 155530 | S17007L |
|  | PE Cy5.5 | CD11c | Invitrogen | 35-0116-42 | 3.9 |

**Supplemental Table 2.** Adipose Immunofluorescence Panel.

| **Primary** | **Company** | **Cat.** | **Clone** |
| --- | --- | --- | --- |
| GSL I Isolectin B4,<br>Biotinylated | Vector<br>Laboratories | B-1205-.5 | N/A |
| Rat anti-mouse RELMa | Invitrogen | MA5-23898 | 226033 |
| Rat anti-mouse Lyve1,<br>eBioscience | Invitrogen | 14-0443-82 | ALY7 |
| Rabbit anti-mouse<br>Serpine1 | Abcam | AB28207 | N/A |
| Rabbit anti-mouse<br>COL6A1 | Proteintech | 17023-1-AP | N/A |
| **Secondary** | **Company** | **Cat.** | **Clone** |
| SAV AF488 | Invitrogen | S32354 | N/A |
| Goat anti-Rat Cy3 | Invitrogen | A10522 | N/A |
| Goat anti-Rabbit Cy5 | KPL | 072-02-15-06 | N/A |

