## Supplementary figures and images for "Helminth infection induces RELMα-dependent adipose tissue transcriptional reprogramming and protection against diet-induced obesity"

### Supplemental Figure 1

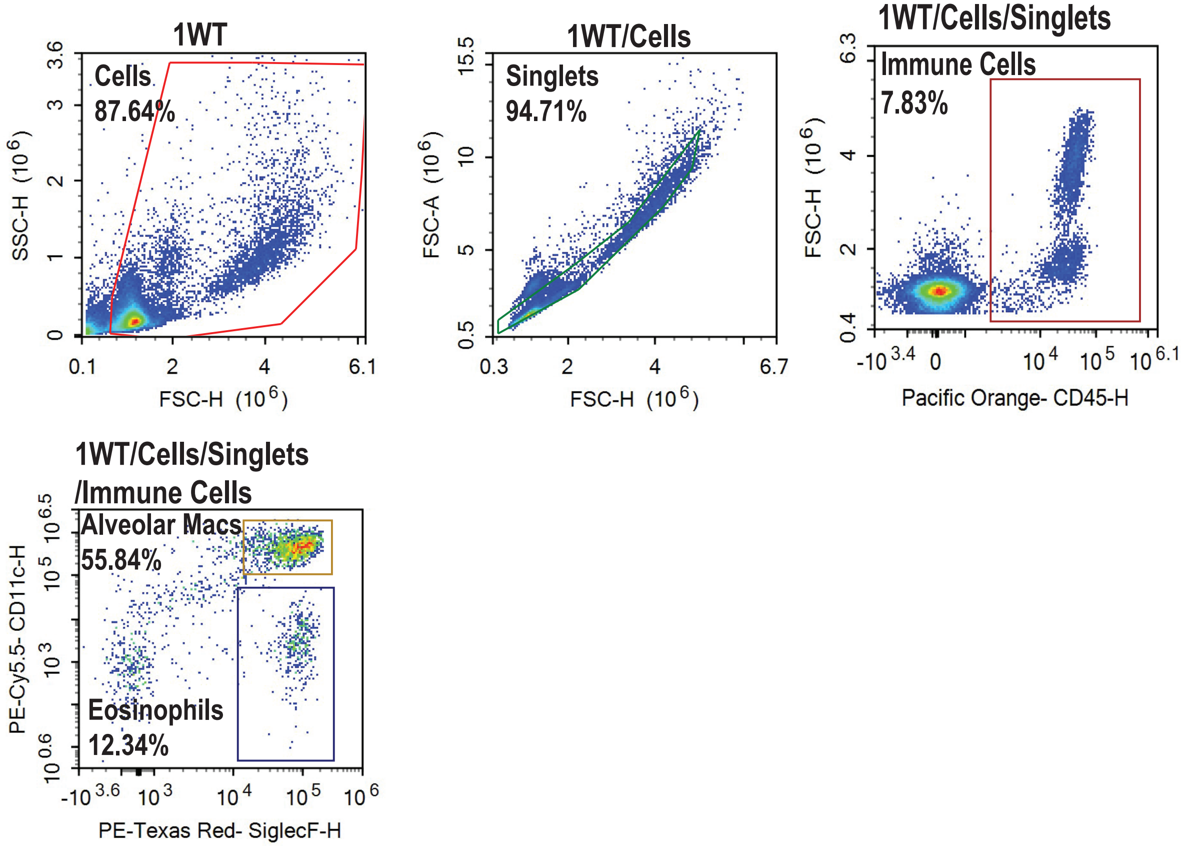

### Supplemental Figure 2

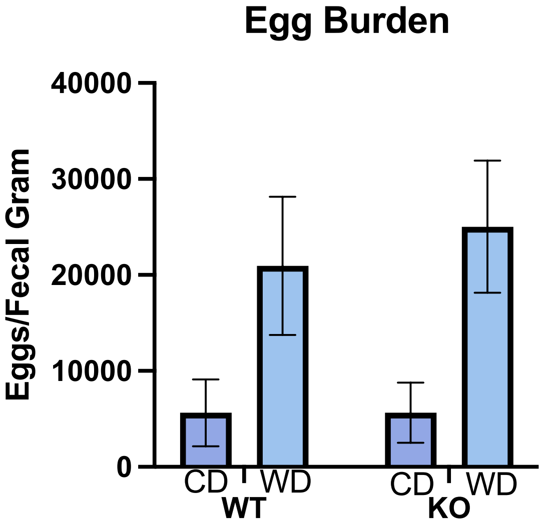

### Supplemental FIgure 3

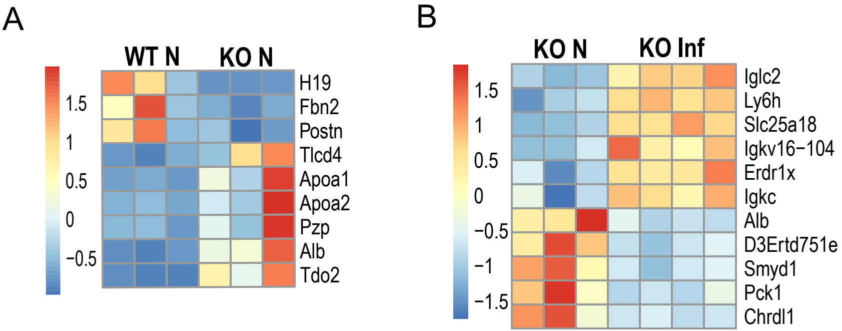
